# Constitutively active mDia1 enables visualization of actin assembly in filopodia

**DOI:** 10.64898/2026.05.27.728329

**Authors:** Brian K. Haarer, Amanda Davenport, Laura M. Haney, Morgan L. Pimm, Alexander D. Nobles, Krishna Patel, Jessica L. Henty-Ridilla

**Affiliations:** Department of Biochemistry & Molecular Biology, SUNY Upstate Medical University, Syracuse, NY, USA; Department of Cell & Developmental Biology, SUNY Upstate Medical University, Syracuse, NY, USA; Department of Neuroscience & Physiology, SUNY Upstate Medical University, Syracuse, NY, USA

**Keywords:** mDia1, formin, actin assembly, filopodia dynamics

## Abstract

Visualizing individual actin filaments in living cells is notoriously difficult. Formins, which track filament plus ends, are powerful probes of actin assembly, yet their behavior in cells is incompletely understood. We dissect the formin mDia1 to reveal how regulatory states control filament assembly. Using TIRF microscopy, we directly compare new regulatory mutants with canonical activity constructs (full-length, FH1-C, FH2-C, and ΔDAD). A two amino-acid substitution mutant, mDia1(CA), is constitutively active and drives strong actin nucleation and elongation. In cells, SNAP-mDia1(CA) is enriched in filopodia, where it forms bright, persistent signals at the tips and dim puncta in the shafts. SNAP-mDia1(CA) increases filopodia number but not elongation rate or length, regardless of its use as an overexpression or rescue construct. Although mDia1(CA) is not a universal actin plus-end tracker in this system, its selective enrichment at filopodial plus ends establishes it as a mechanistically distinct probe for interrogating filopodial actin dynamics.

## Introduction

Direct visualization of actin filament assembly in living cells remains a central challenge in cytoskeletal biology. In the microtubule field, fluorescently tagged plus-end-binding proteins provide reliable fiducial markers that report polymer growth with high temporal resolution and enable quantitative tracking of individual polymers within complex cellular environments (Morrison et al., 2002; Stepanova et al., 2003; Slep and Vale, 2007). No comparable genetically encoded system exists for actin filaments. Current approaches including fluorescent speckle microscopy, FRAP, photoconversion, PALM, or live-cell PAINT, primarily report bulk filament turnover or network behavior (Betzig et al., 2006; Bhaskar et al., 2025; Lacy et al., 2019; Waterman-Storer and Salmon, 1998). Although these methods have enabled important insights into filament assembly and disassembly in cells, tracking individual actin filaments becomes difficult due to network density and resolution limits. In addition, many fluorescent probes perturb actin dynamics, necessitating minimally invasive labels that often reduce spatial or temporal resolution (Lemieux et al., 2014; Melak et al., 2017). Thus, direct observation of defined actin filament plus-end behaviors in cells remains limited.

Formin proteins are attractive candidates for addressing this gap. These processive actin assembly factors associate with actin filament plus ends, nucleate filaments, and often promote accelerated elongation with profilin (Pruyne et al., 2002; Sagot et al., 2002; Romero et al., 2004; Kovar et al., 2006). Given the high abundance of profilin-actin in cells, formins such as mDia1 may be uniquely positioned among actin regulatory proteins to sustain productive actin filament plus-end elongation through their ability to directly recruit and utilize profilin-bound actin monomers (Tang et al., 2025; Suarez et al., 2015; Rotty et al., 2015). Mammalian cells express multiple formins with diverse regulatory and structural properties that include filament nucleation, elongation, capping, bundling, severing, and microtubule association (Schönichen and Geyer, 2010; Valencia and Quinlan, 2021). Among them, the Diaphanous-related formin, mDia1, is broadly expressed and has been implicated in filopodia formation (Higashida et al., 2004; Goh and Ahmed, 2012), stress fiber assembly (Hotulainen and Lappalainen, 2006), and actin-microtubule interactions (Ishizaki et al., 2001; Roth-Johnson et al., 2014; Palazzo et al., 2001; Pimm and Henty-Ridilla, 2021). Importantly, mDia1 promotes filament nucleation and elongation without intrinsic bundling activity, making it a useful candidate for marking sites of actin assembly without directly reorganizing filament architecture via crosslinking.

mDia1 actin assembly activities are controlled by intramolecular autoinhibition mediated by interactions between its Diaphanous inhibitory domain (DID) and C-terminal autoregulatory (DAD) region (Li and Higgs, 2003; Rose et al., 2005; Lammers et al., 2005; Nezami et al., 2006; Otomo et al., 2010). Relief of this interaction through deletion (Li and Higgs, 2005), mutation (Lammers et al., 2005; Lakha et al., 2021), or competitive binding interactions with regulatory factors (i.e., RhoA, IQGAP1) activates filament assembly (Lammers et al., 2005; Otomo et al., 2005; Higashi et al., 2008; Brandt et al., 2007; Pimm et al., 2024). Although prior studies have examined specific mDia1 mutants or regulatory interactions in vitro and in cells (Lammers et al., 2005; Seth et al., 2006; Copeland et al., 2007) and related regulatory principles have been explored more extensively for the closely related isoform mDia2 (Block et al., 2008; Barzik et al., 2014), these analyses have largely been conducted in isolation. A systematic, side-by-side comparison directly linking defined biochemical properties of mDia1 variants to their behavior in living cells is still lacking.

Here, we dissect mDia1 regulatory and structural domains to determine whether defined constructs can function as reporters of actin assembly in cells. Using Total Internal Reflection Fluorescence (TIRF) microscopy, we validate the nucleation and elongation activities of full-length, truncated, and constitutively active mDia1 proteins relative to well-established formin constructs. We then perform advanced imaging of SNAP-tagged proteins in Neuroblastoma-2A (N2A) cells, including mDia1 knockout lines, to assess how these variants shape filament dynamics in live cells. The constitutively active formin, SNAP-mDia1(CA), does not label all actin plus ends and does not function as a universal actin plus-end tracker. Instead, SNAP-mDia1(CA) selectively marks a defined subset of plus ends engaged by mDia1 and is competent for formin-mediated assembly. In cells, SNAP-mDia1(CA) is enriched at filopodia, where it marks discrete assembly sites. Quantitative analysis reveals two distinct populations of SNAP-mDia1(CA) puncta within filopodia, including bright accumulations at tips and dim puncta along filopodia shafts. Despite an increase in filopodia number upon mDia1 activation, elongation rates and steady-state lengths remain largely unchanged. Thus, SNAP-mDia1(CA) highlights mDia1-mediated filament assembly in filopodia, providing a tool to directly observe formin activity in cells.

## Results

### Structure-guided activation of mDia1 preserves actin assembly activities

To determine whether mDia1 could serve as a reporter of actin assembly states in cells, we systematically truncated mDia1 to explore how discrete domains contribute to actin assembly in vitro. We generated and purified a panel of mDia1 proteins spanning key regulatory and catalytic regions (Figure 1A and Supplementary Figure 1A-C). These included full-length mDia1 (FL; residues 1-1255); an N-terminal fragment lacking canonical actin assembly domains but retaining the GTPase-binding domain (GBD) required for RhoA activation and the basic region implicated in membrane localization (NT; residues 1-570) (Li and Higgs, 2005; Ramalingam et al., 2010); a truncation lacking the armadillo repeat region (ΔARR; residues 257-1255); a DAD deletion that relieves autoinhibition (ΔDAD; 1-1175) (Li and Higgs, 2005; Pimm et al., 2024); a formin homology (FH)1 to C-terminal construct competent for profilin-dependent elongation (FH1-C; residues 571-1255) (Kovar et al., 2006); and a FH2 to C-terminal construct previously shown to nucleate and cap actin filaments (FH2-C; residues 739-1255) (Li and Higgs, 2003). We also engineered a mutant in the full-length protein predicted to be constitutively active (mDia1(CA)), containing A256D and I259D substitutions within the Diaphanous Inhibitory Domain (DID) (Figure 1B) (Lammers et al., 2005; Rose et al., 2005; Block et al., 2008; Nezami et al., 2006). These substitutions were designed to disrupt the autoinhibitory interaction between the DID-DAD interface while preserving the domain architecture of FL-mDia1. This design enabled direct comparison between an activated full-length formin and commonly used truncation-based proteins competent for actin assembly.

**Figure 1.**
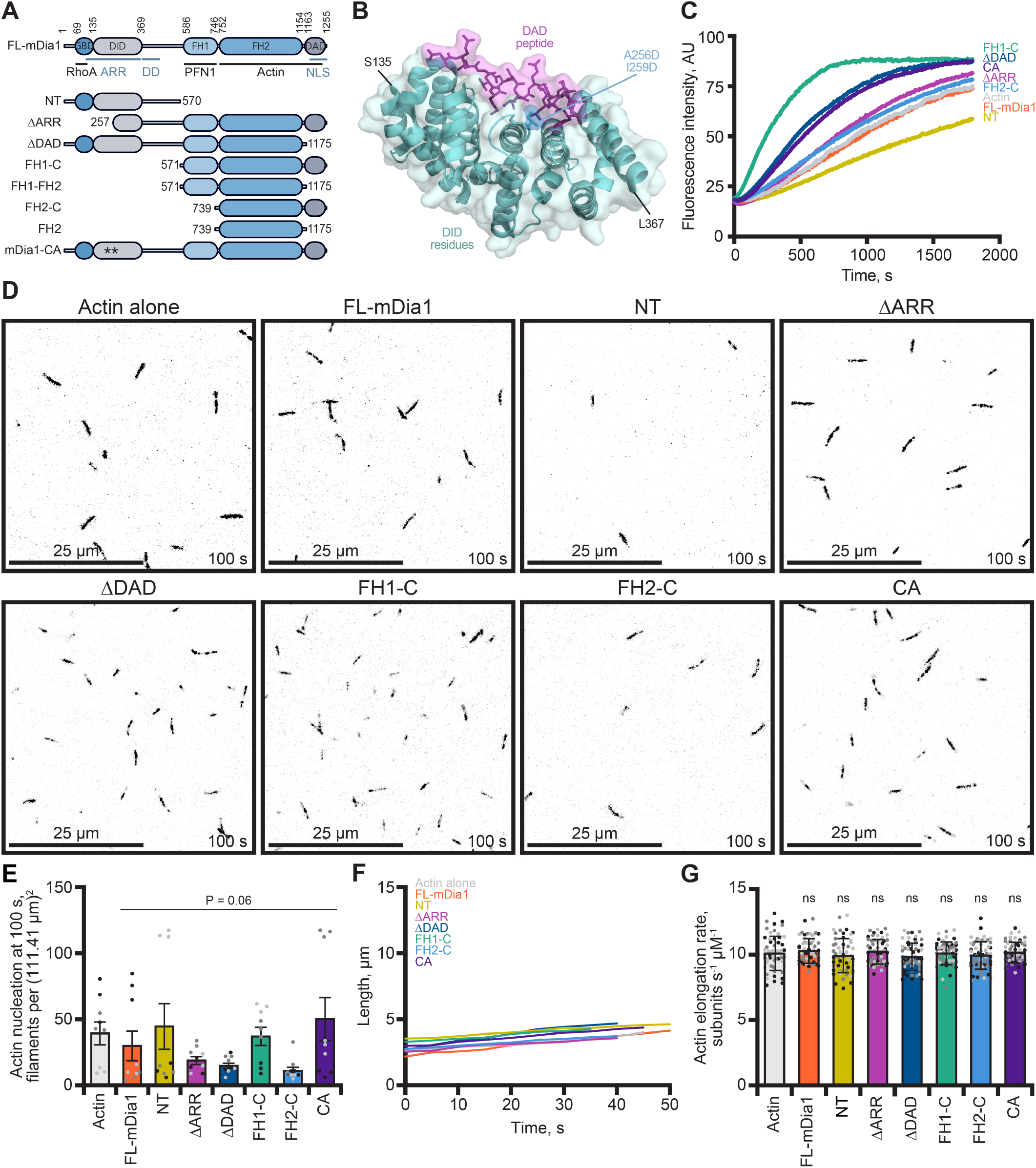
Diaphanous Inhibitory Domain (DID) mutations convert full-length mDia1 into a nucleation-competent formin. **(A)** Schematic of mDia1 domain organization and constructs used in this study. Domains include the GTPase-binding domain (GBD), diaphanous inhibitory domain (DID), armadillo repeat region (ARR), formin homology domains FH1 and FH2, and the diaphanous autoregulatory domain (DAD). Constructs are: full-length (FL), N-terminal fragment (NT), ΓARR, ΓDAD, FH1-C, FH2-C, and constitutively active mDia1(CA), which contains A256D and I259D substitutions within the DID (asterisks). **(B)** Structural model of the DID–DAD autoinhibitory interface (PDB 2F31; Nezami et al., 2006). The DID is shown as a cartoon (teal) and the DAD peptide as sticks (magenta). Residues mutated in mDia1(CA) (A256, I259) are shown in blue. **(C)** Bulk pyrene-actin polymerization assays containing 2 µM actin (5% pyrene-labeled) in the absence or presence of the indicated mDia1 constructs in (A). Traces are representative of N = 3 independent experiments. **(D)** TIRF microscopy images of 1 µM actin (10% Alexa Fluor 488 Lys-label) polymerizing alone or in the presence of 1 nM indicated mDia1 constructs, acquired 100 s after initiation of polymerization. Scale bars, 25 µm. **(E)** Quantification of filament nucleation from experiments performed with 1 µM actin (10% Alexa Fluor 488 Cys374-label), expressed as the number of filaments per field of view (FOV) at 100 s from initiation (N = 3 independent experiments, indicated by shading). Each dot represents the number of filaments per FOV. Statistical analysis was performed by one-way ANOVA, compared to actin alone control; ns, not significant (P > 0.05). **(F)** Representative length-versus-time plots of individual filaments from reactions shown in (D). **(G)** Mean filament elongation rates calculated from TIRF assays. Each dot represents one filament (n = 54 filaments per condition); bars are means ± SD; shading denotes independent experiments (N = 3 reactions per condition). Statistical analysis was performed by one-way ANOVA, compared to actin alone control; ns, not significant (P > 0.05).

### mDia1(CA) relieves autoinhibition to promote actin filament nucleation

We first assessed actin assembly of mDia1 proteins using bulk pyrene polymerization assays and TIRF microscopy. In pyrene actin assembly assays (Figure 1C), constructs lacking canonical autoinhibitory constraints (i.e., FH1-C, FH2-C, and ΔDAD) enhanced filament polymerization relative to actin alone controls. Full-length (FL) mDia1 exhibited minimal activity, consistent with autoinhibition. Notably, mDia1(CA) increased bulk polymerization relative to the full-length protein and to levels approaching FH1-C, indicating that the A256D/I259D substitution at least partially relieves autoinhibition in the context of the full-length protein.

To directly quantify filament nucleation, we performed TIRF microscopy using 1 µM actin and 1 nM mDia1 constructs (Figures 1D-E and Movie 1). At 100 s after initiation of polymerization, actin alone reactions contained an average of 41.4 ± 8.5 filaments per field of view (± SEM). No formin treatment significantly elevated filament nucleation compared to actin alone control (P = 0.06), including the autoinhibited FL-mDia1 which exhibited minimal nucleation activity (31.2 ± 11.2), the N-terminal fragment (45.9 ± 17.3), and FH1-C (38.4 ± 6.9). ΔARR, ΔDAD, and FH2-C lacked significant assembly activity compared to actin alone or FL-mDia1, likely due to intact filament capping capacity (20.1 ± 2.9, 16 ± 2.1, and 12.2 ± 2.8, respectively; P = 0.47). mDia1(CA) had similar nucleation counts compared to actin alone controls (51.6 ± 16.4; P > 0.99).

Despite differences in filament number, elongation rates were largely unchanged across constructs (Figure 1F-G; P = 0.25), indicating that these perturbations primarily affect nucleation competence rather than basal filament elongation. Together, these findings demonstrate that targeted disruption of the DID-DAD interface is sufficient to convert FL-mDia1 into a nucleation-competent protein while preserving its native domain organization.

### mDia1(CA) supports profilin-dependent, processive filament elongation

Accelerated actin filament elongation in the presence of profilin is a defining feature of processive formin activity at actin filament plus ends. Thus, we tested whether mDia1 constructs could support profilin-dependent actin assembly using both bulk pyrene assays and single-filament TIRF microscopy. In pyrene assays performed with 5 µM profilin-1 (PFN1), active formin fragments (FH1-C, FH2-C, and ΔDAD) robustly enhanced actin assembly relative to actin-profilin controls (Figure 2A-B), consistent with previous studies (Li and Higgs, 2003; Kovar et al., 2006; Paul et al., 2008; Maiti et al., 2012). In contrast, full-length mDia1 and the N-terminal fragment (NT) did not increase polymerization, consistent with autoinhibition and absence of FH1-mediated profilin interactions, respectively. Importantly, mDia1(CA) supported profilin-dependent actin assembly at levels approaching FH1-C, indicating that disruption of the DID-DAD interface induces functional activation of the full-length protein.

**Figure 2.**
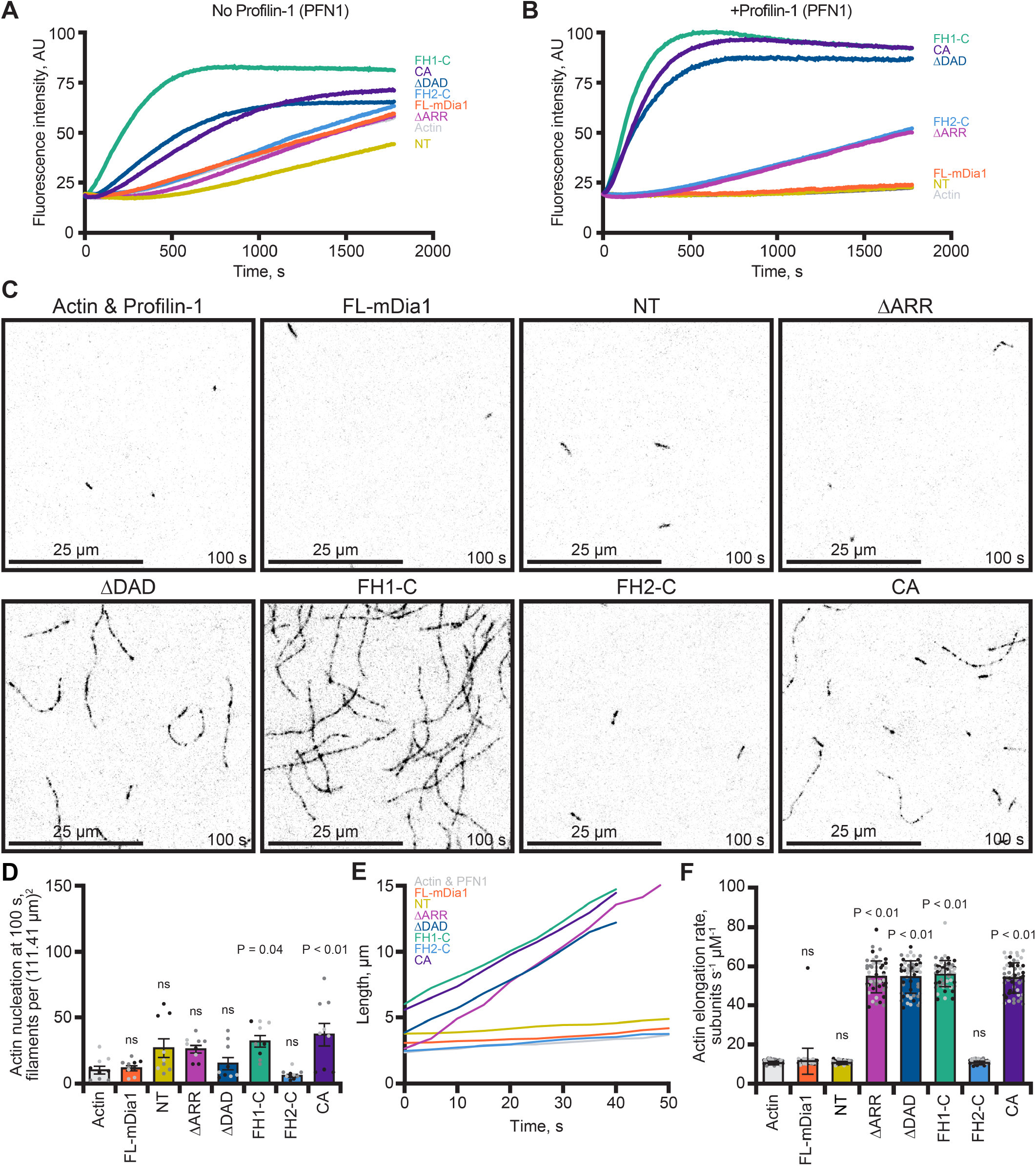
mDia1(CA) enables accelerated formin elongation in the presence of profilin-1 (PFN1). **(A)** Bulk pyrene-actin polymerization assays containing 2 µM actin (5% pyrene-labeled) and 1 nM of the indicated mDia1 constructs in the absence of PFN1. **(B)** Bulk polymerization assays as in (A), performed in the presence of 5 µM PFN1. Traces are representative of N = 3 independent experiments. **(C)** TIRF images acquired 100 s after initiation of polymerization from reactions containing 1 µM actin (10% Alexa Fluor 488 Lys-label), 5 µM PFN1, and 1 nM of the indicated mDia1 constructs. Scale bars, 25 µm. **(D)** Quantification of filament nucleation from experiments performed with 1 µM actin (10% Alexa Fluor 488 Cys374-label), expressed as the number of filaments per FOV at 100 s from initiation (N = 3 independent experiments, indicated by shading). Each dot represents the filament count from an FOV. Statistical significance compared to actin and PFN1 control; ns, not significant (P > 0.05). **(E)** Representative length-versus-time plots of individual filaments from reactions in (C), illustrating accelerated elongation in the presence of active mDia1 constructs. **(F)** Mean filament elongation rates calculated from TIRF assays. Each dot represents a single filament (n = 54 filaments per condition); bars are means ± SD shading denotes independent experiments (N = 3 reactions per condition). Statistical analysis was performed by one-way ANOVA, compared to actin and PFN1 control; ns, not significant (P > 0.05).

To determine whether the inactivity of full-length mDia1 reflected autoinhibition rather than protein misfolding, we added recombinant RhoA to reactions containing FL-mDia1 or ΔARR in the presence of PFN1 (Supplementary Figure 1D-E). Addition of RhoA significantly enhanced actin assembly by FL-mDia1, consistent with canonical disruption of DID-DAD autoinhibition (Li and Higgs, 2003, 2005; Otomo et al., 2005; Maiti et al., 2012). In contrast, RhoA did not further stimulate ΔARR, which lacks the GTPase-binding region required for RhoA-mediated activation (Figure 1A). These results demonstrate that purified FL-mDia1 remains functional but autoinhibited under these conditions.

We next directly quantified filament nucleation and elongation rates using single-filament TIRF (Figure 2C-F and Movie 2). As expected, the addition of profilin to TIRF reactions containing polymerizing actin filaments reduced the overall number of filaments from 41.4 ± 8.5 (Figure 1E) to 9.7 ± 2.8 filaments per FOV (Figure 2C-D; P = 0.27). Similar to conditions without profilin, most formin treatments did not significantly change filament nucleation compared to the actin and profilin control. The exceptions were the most potent filament nucleators, FH1-C (31.1 ± 4.3; P = 0.04) and mDia1(CA) (36.2 ± 8.4; P < 0.01). In the absence of profilin, elongation rates were comparable across all constructs and indistinguishable from actin-alone controls that elongated on average 10.1 ± 1.3 subunits s^-1^ µM^-1^ ± SD (Figure 1F-G; P = 0.254), indicating that mDia1 constructs do not substantially alter plus-end elongation under these conditions. In contrast, addition of 5 µM profilin-1 (PFN1) revealed pronounced construct-dependent differences in elongation behavior (Figure 2C-F, Supplementary Figure 1F-G, and Movie 2). FH1-C, ΔDAD, and ΔARR significantly increased mean elongation rates to 55.2 ± 6.8 (P < 0.01), 53.9 ± 8.4 (P < 0.01), and 54.0 ± 8.2 subunits s^-1^ µM^-1^ ± SD (P <0.01), respectively. This is consistent with profilin-mediated delivery of actin monomers to processively associated formin at filament plus ends (Kovar et al., 2006; Romero et al., 2004; Courtemanche and Henty-Ridilla, 2024). Similar to the pyrene assays above, full-length mDia1 and NT did not significantly alter elongation rates. Notably, mDia1(CA) supported profilin-dependent elongation at levels consistent with FH1-C (CA: 53.4 ± 7.9 subunits s^-1^ µM^-1^ ± SD; P = 0.74, compared to FH1-C and PFN1 condition), demonstrating that disruption of the DID-DAD interface restores efficient coupling between profilin-actin delivery and formin-mediated plus-end elongation in the full-length protein.

Fast-growing filaments assembled in the presence of profilin frequently exhibited reduced fluorescence intensity, a qualitative feature previously associated with processively elongating formin-bound filaments under similar (Cys374-label) conditions (Kovar et al., 2006). We therefore quantified the fraction of filaments exhibiting dim rapidly elongating growth segments at 200 s after reaction initiation (Supplementary Figure 1F-G and Movie 3). This dim-fast filament population was most prominent in reactions containing active FH1-domain constructs, including FH1-C, ΔDAD, or mDia1(CA), whereas FL-mDia1 or NT did not display this behavior. We interpret these dim, rapidly elongating filaments as consistent with sustained formin engagement during profilin-dependent actin filament plus-end elongation.

Thus, because rapidly elongating filaments assembled from Cys374-labeled actin exhibited reduced fluorescence intensity in the presence of profilin, representative images in Figure 2C were acquired using lysine-labeled actin to improve filament visualization. Quantitative analyses of elongation behavior and dim-filament frequency were performed using Cys374-labeled actin reactions, where the dim-fast phenotype was most readily resolved. Together, these results demonstrate that mDia1(CA) restores profilin-dependent, processive elongation behavior characteristic of active FH1-FH2-containing formin constructs while preserving the native full-length regulatory architecture.

### Activated SNAP-mDia1 constructs form filopodiaassociated puncta and promote filopodia formation in N2A cells

To determine whether activated mDia1 could be used to monitor formin-associated actin assembly in cells, we generated N-terminally SNAP-tagged versions of the constructs characterized in Figures 1-2 and transiently expressed them in wild-type mouse Neuroblastoma-2A (N2A) cells (Figure 3) or mDia1 knockout N2A cells (Figures S2 and S3). SNAP labeling was performed using empirically optimized low concentrations of SNAP-Cell 647-SiR to resolve discrete mDia1-positive structures (Pimm et al., 2022). Constructs were co-expressed with GFP-actin to visualize actin networks (Figure 3A). Western blot analysis demonstrated expression levels were comparable to endogenous mDia1 (Supplementary Figure 2A-D), and localization patterns were similar in wild-type and knockout backgrounds (Figure 3A; Supplementary Figure 3A), indicating the phenotypes described below do not require endogenous mDia1 and are unlikely to arise from gross overexpression artifacts.

**Figure 3.**
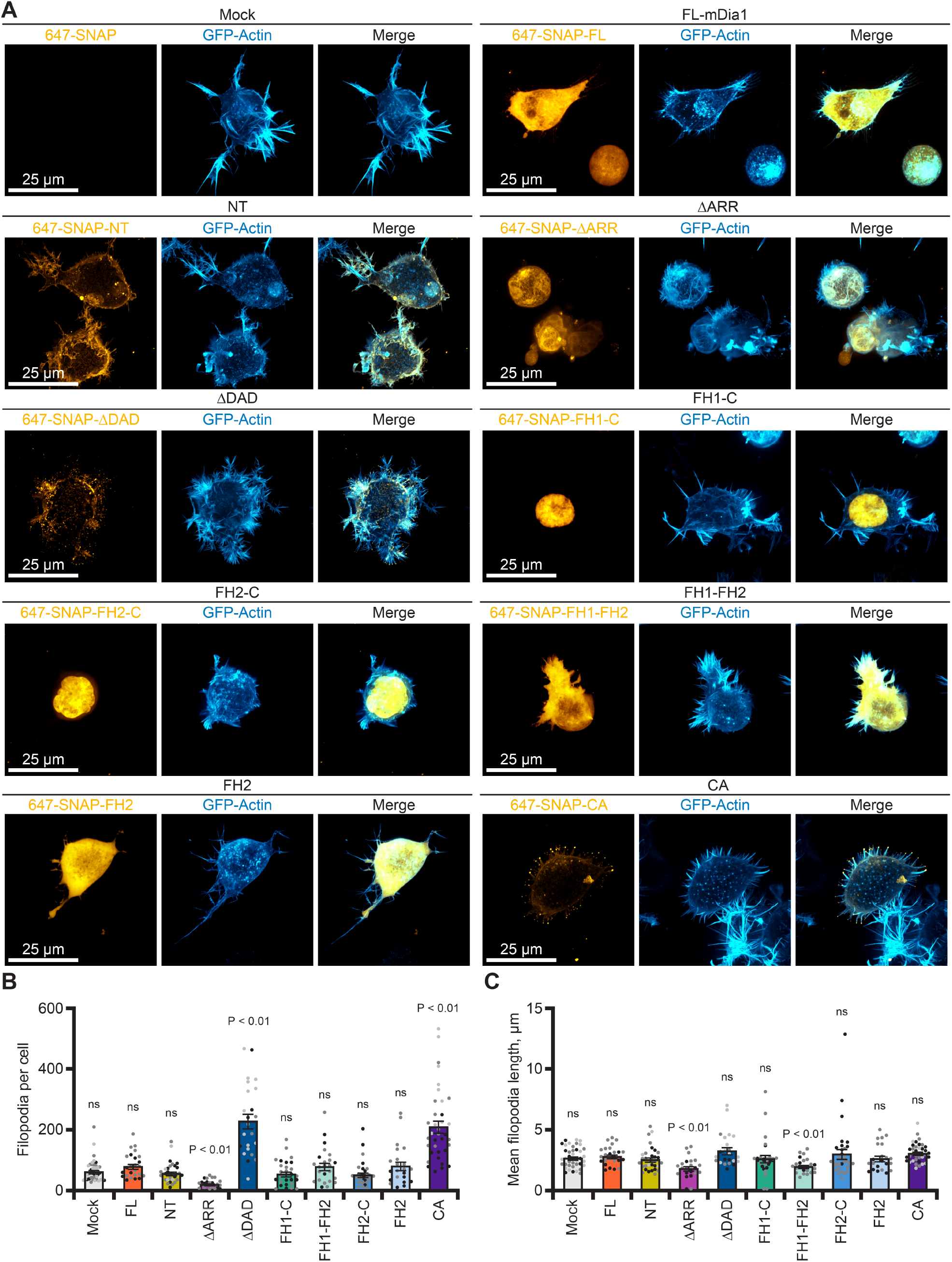
SNAP-mDia1(CA) is enriched at filopodia tips and increases their number. **(A)** Live-cell confocal images of wild-type N2A cells expressing the indicated N-terminally SNAP-tagged mDia1 constructs. SNAP signal (gold), GFP-actin (blue), and merged images are shown. Maximum intensity projections are displayed. Scale bars, 25 µm. **(B)** Quantification of filopodia number per cell from 3D image stacks as in (A). Each dot represents one cell (n = 22-40 cells per condition); shading denotes N = 2-4 independent experiments (separate transfections performed on different days). Statistical analysis was performed by one-way ANOVA, compared to mock; ns, not significant (P > 0.05). **(C)** Average length of filopodia per cell from datasets as in (A). Each dot represents the mean filopodial length for a single cell (n = 22-40 cells per condition). Shading denotes N = 2-4 independent experiments. Statistical analysis was performed by one-way ANOVA, compared to mock; ns, not significant (P > 0.05).

SNAP-FL-mDia1 was diffuse throughout the cytoplasm and showed no obvious enrichment at actin-rich structures (Figure 3A), consistent with autoinhibited behavior observed in vitro (Figures 1 and 2). In contrast, SNAP-mDia1(NT) localized prominently to the cell cortex and plasma membrane, consistent with established membrane-targeting functions within the N-terminus (Seth et al., 2006; Rousso et al., 2013), but lacked detectable enrichment at filopodia tips. Interestingly, constructs that were highly active in vitro were not necessarily associated with actin-based structures in cells. SNAP-mDia1(FH1-C) and SNAP-mDia1(FH2-C) accumulated predominantly within the nucleus despite robust biochemical activity in vitro. This is consistent with exposure of the previously described C-terminal nuclear localization sequence upon removal of upstream regulator domains (Copeland et al., 2007). Removal of the DAD region from these truncations (i.e., SNAP-mDia1-FH1FH2 or SNAP-mDia1-FH2) reduced nuclear enrichment and produced a more diffuse cytoplasmic localization (Figure 3A and Supplementary Figure 3A). Together, these observations indicate that catalytic activity alone is insufficient for productive localization to cellular actin structures and suggest that retention of native regulatory architecture contributes to appropriate spatial targeting in cells.

In contrast, SNAP-mDia1(ΔDAD) and SNAP-mDia1(CA) formed discrete cytoplasmic puncta that were frequently enriched along filopodia and at filopodia tips (Figure 3A and Supplementary 4A). These puncta were spatially restricted and comparatively persistent. Because mDia1 has been implicated in actin-microtubule crosstalk (Gaillard et al., 2011; Kato et al., 2001; Daou et al., 2014; Bartolini et al., 2016), we tested whether expression of SNAP-mDia1(CA) altered microtubule polymerization dynamics. Expression of SNAP-mDia1 constructs did not reveal conspicuously altered microtubule organization (Supplementary Figure 4A). Further, quantification of EB1 comets revealed no significant differences in microtubule velocity or comet number relative to mock-transfected cells (Supplementary Figure 4B-D), indicating that the observed puncta and filopodial phenotypes are unlikely to result from secondary effects on microtubule polymerization.

**Figure 4.**
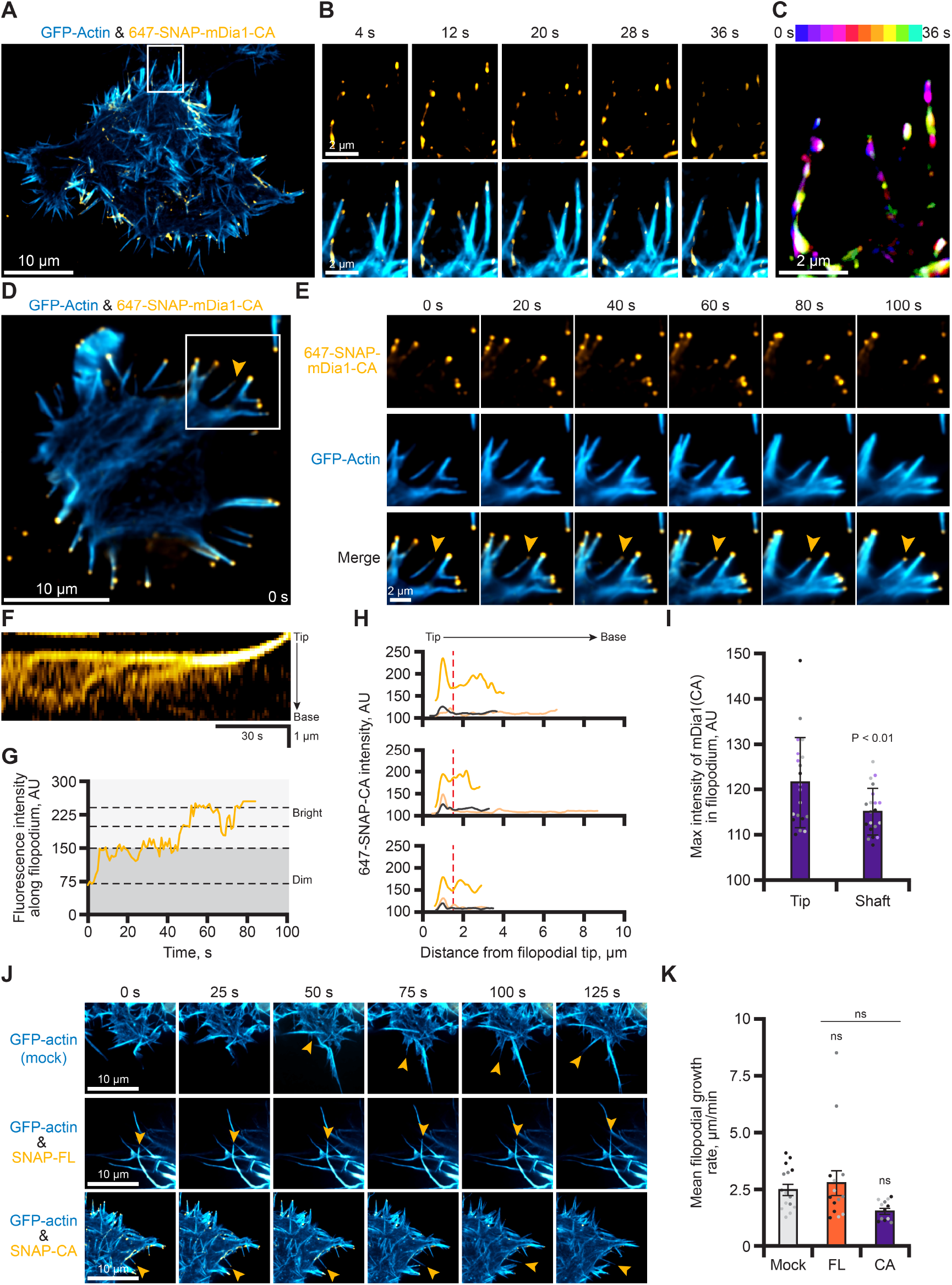
SNAP-mDia1(CA) forms distinct bright and dim puncta populations within filopodia without altering extension rates. **(A)** Representative SoRa confocal image of a live N2A cell co-expressing GFP-actin (blue) and SNAP-mDia1(CA) (gold), showing heterogeneous SNAP signal intensities within filopodia. Scale bar, 10 µm. **(B)** Magnified time-lapse montage of the region shown in (A), highlighting filopodia containing bright and dim SNAP-mDia1(CA) puncta over 36 s. Scale bars, 2 µm. **(C)** Time-encoded projection of the 647-SNAP signal in the region highlighted in (A-B), spanning 36 s. Frames are color-coded by time; greater color variation indicates dynamic signal, whereas white indicates minimal or no movement over the acquisition period. **(D)** Representative TIRF image of a live N2A cell co-expressing GFP-actin (blue) and SNAP-mDia1(CA) (gold). Arrow highlights an individual filopodium in which SNAP signal accumulates at the tip over time. **(E)** Magnified time-lapse montage of the filopodium shown in (D), illustrating progressive SNAP-mDia1(CA) enrichment at the tip over 100 s. Scale bar, 2 µm. **(F)** Kymograph of the filopodium shown in (E), showing accumulation of SNAP-mDia1(CA) signal at the tip over time. Diagonal trajectories indicate processive movement of puncta toward the tip. Time scale, 30 s. Distance scale, 1 µm. **(G)** Line scan derived from the kymograph in (F), showing increasing SNAP-mDia1(CA) intensity at the filopodial tip as puncta accumulate over time. **(H)** Line scans from multiple filopodia (different colors) aligned at the tip, showing SNAP-mDia1(CA) intensity profiles along filopodial length. The red dotted line marks the boundary between the tip region (0-1.5 µm from the tip) and the shaft (>1.5 µm from the tip). Graphs represent examples from independent replicates. **(I)** Quantification of maximum SNAP-mDia1(CA) puncta intensity in tip (0-1.5 µm) and shaft (1.5-3 µm) of filopodia. Each point represents values from a filopodium (n = 21 filopodia per condition) from N = 3 independent transfections (shading). Statistical analysis was determined by one-way t-test, comparing tip and shaft intensities. **(J)** Time-lapse SoRa confocal montage of growing filopodia in wild-type N2A cells expressing GFP-actin (mock; blue) or GFP-actin and SNAP-mDia1(FL) or SNAP-mDia1(CA) in gold. Scale bars, 10 µm. **(K)** Quantification of filopodial elongation rates in cells expressing the indicated SNAP-mDia1 constructs. Each dot represents the mean elongation rate per filopodium; shading denotes independent cells and transfections. Statistical analysis was performed by one-way ANOVA, compared to mock; ns, not significant (P > 0.05).

Expression of SNAP-mDia1(ΔDAD) or SNAP-mDia1(CA) significantly increased the number of filopodia per cell (Figures 3B and Supplementary Figure 3B), whereas average filopodial length was unchanged or only modestly affected (Figures 3C and Supplementary Figure 3C). Notably, SNAP-mDia1(CA) produced filopodia phenotypes comparable to the ΔDAD construct while preserving the full-length domain organization of mDia1. Similar localization and filopodial phenotypes were observed in mDia1 knockout cells (Supplementary Figure 3), indicating that these behaviors do not depend on endogenous mDia1. Thus, these results identify mDia1(CA) as an activated FL-mDia1 variant that retains appropriate subcellular targeting while forming discrete puncta enriched at filopodia tips and promoting filopodia formation in cells.

### Distinct SNAP-mDia1(CA) puncta populations within filopodia

To define how activated SNAP-mDia1 behaves within filopodia, we performed live-cell imaging of SNAP-mDia1(CA) and GFP-actin in N2A cells using spinning disk confocal and TIRF microscopy (Figure 4). Live time-lapse imaging revealed heterogeneous SNAP-mDia1(CA) puncta within filopodia that differed in both fluorescence intensity and dynamic behavior. Live confocal imaging showed prominent SNAP-mDia1(CA) enrichment at the distal tips of filopodia (Figure 4A-C and Movie 4). Temporal color projections over a 36 s interval further showed that many tip-associated puncta remained somewhat spatially constricted over time, appearing persistent relative to more variable and dynamic puncta observed along filopodia shafts (Figure 4C).

We next examined these populations of puncta using TIRF microscopy to better resolve SNAP-mDia1(CA) behavior at the cell cortex (Figure 4D-G and Movie 5). Time-lapse imaging revealed progressive accumulation of SNAP-mDia1(CA) signal at a subset of filopodia tips (Figure 4D-E). Notably, kymograph analysis of the filopodium highlighted in Figure 4E demonstrated that bright tip-associated puncta remained persistently localized at the filopodia tips over time (Figure 4F). Quantification of fluorescence intensity from the kymograph further revealed transitions between discrete dim and bright SNAP-mDia1(CA) states, in which dimmer shaft-associated puncta underwent sustained accumulation at the filopodial tip, resulting in progressive tip enrichment of SNAP-mDia1(CA) signal (Figure 4G). Consistent with this observation, line-scan analysis across multiple filopodia showed enrichment of SNAP-mDia1(CA) fluorescence at the tip relative to the shaft region (Figure 4H). Quantification confirmed that maximum puncta intensity was significantly higher at filopodial tips than within the proximal 3 µm of the shaft (Figure 4I; P < 0.01, t-test), indicating that tip-associated puncta represent a spatially distinct population.

In addition to these bright tip puncta, dimmer SNAP-mDia1(CA) puncta were frequently detected along filopodial shafts. These shaft-associated puncta were typically more dynamic than tip-associated structures and occasionally moved with velocities similar to measured filopodial extension rates. Although these observations may be associated with actin assembly within filopodia, expression of SNAP-mDia1(CA) did not significantly alter overall filopodial extension rates relative to mock or SNAP-FL-mDia1 controls (Figure 4J-K; P = 0.06). Together, these data identify at least two spatially and dynamically distinct SNAP-mDia1(CA) puncta populations within filopodia: bright, persistent puncta enriched at tips and dimmer, more dynamic puncta distributed along filopodial shafts. The persistent localization of bright puncta at extending filopodia tips, particularly evident in kymograph analyses, supports the idea that activated mDia1 associates with discrete actin assembly sites within filopodia, while preserving overall filopodia growth dynamics.

## Discussion

Understanding where and when actin filaments assemble in cells remains a central challenge in cell biology because filament nucleation and polymerization are tightly coupled to local signaling and mechanical cues. Formins are attractive candidates for visualizing subsets of active actin assembly because they processively associate with filament plus ends downstream of Rho-family GTPase signaling. However, the details underlying how distinct regulatory and catalytic regions of formins influence productive actin engagement in cells has remained incompletely understood. Here, we systematically analyzed a panel of mDia1 proteins in vitro and in living cells to determine whether an activated full-length mDia1 could be used to visualize spatially restricted actin assembly behaviors.

Our findings demonstrate that relief of autoinhibition within the context of FL-mDia1 is sufficient to restore both nucleation and profilin-dependent elongation activity while preserving productive cellular localization. In contrast, many truncated constructs that retained robust biochemical activity failed to recapitulate normal subcellular behavior in cells. In particular, FH1-FH2-containing truncations accumulated predominantly within the nucleus, consistent with previous observations that removal of upstream regulatory regions exposes a C-terminal nuclear localization sequence (Copeland et al., 2007; Gould et al., 2011; Vizcarra et al., 2014). These observations establish an important principle: catalytic competence alone does not predict productive actin assembly behavior in cells. Instead, regulatory and targeting domains play essential roles in determining where formins can engage actin networks.

Activated SNAP-mDia1(CA) localized prominently to filopodia, where it formed discrete puncta enriched at tips and along filopodial shafts. Live-cell imaging further revealed at least two spatially and dynamically distinct puncta populations within filopodia. Bright puncta accumulated at tips and remained comparatively persistent once established, whereas dimmer shaft-associated puncta were more dynamic and transient. Although the dim puncta frequently appeared within actively remodeling filopodia, their rapid movements and comparatively low fluorescence intensity limited reliable tracking over extended intervals. Consequently, we cannot directly conclude that these puncta correspond to processively elongating plus ends. Importantly, however, overall filopodia extension rates were not significantly altered by SNAP-mDia1(CA) expression, indicating that activated mDia1 increases filopodia number without globally accelerating protrusion dynamics. Together, these observations suggest that activated mDia1 associates with spatially restricted actin assembly states within filopodia rather than uniformly enhancing protrusive growth. More broadly, these findings highlight that formin behavior in cells is shaped not only by intrinsic polymerization activity, but also by local regulatory context, membrane association, and subcellular organization. Notably, we rarely observed comet-like movements outside filopodia, indicating that cytoplasmic plus-end tracking is not a dominant behavior for mDia1 in N2A cells and emphasizing that formin dynamics differ substantially from simplified analogies to microtubule plus-end tracking proteins.

Our findings also support the broader idea that productive actin assembly in cells may depend on selective access to profilin-bound actin monomer pools (Suarez et al., 2015; Rotty et al., 2015; Tang et al., 2025). Formins are uniquely positioned to exploit these pools through FH1-mediated profilin-actin delivery, potentially enabling localized filament elongation downstream of signaling events. In this context, the restricted localization and behavior of SNAP-mDia1(CA) suggest that productive formin engagement is not solely determined by catalytic activation, but also by whether local cellular environments support sustained utilization of profilin-actin during assembly (Tang et al., 2025). Although our study does not directly measure monomer partitioning, these observations are consistent with the emerging view that distinct actin regulators compete for or preferentially access different actin assembly states within cells.

By preserving the full-length domain architecture while relieving autoinhibition, SNAP-mDia1(CA) provides a means to visualize a spatially restricted subset of mDia1-associated actin assembly events in cells. Importantly, this probe does not label all sites of filament growth, underscoring why universal markers of actin nucleation and elongation have remained difficult to establish. These findings reinforce the idea that actin polymerization in cells is tightly coupled to regulatory networks that spatially and temporally constrain filament assembly. More broadly, our findings illustrate a general principle for cytoskeletal probes: effective visualization of polymer dynamics requires preservation of both catalytic activity and the local regulatory landscape in cells. Together, this work establishes activated full-length mDia1 as a useful platform for probing localized formin-associated actin assembly and provides a framework for dissecting how RhoA signaling, membrane organization, and mechanical inputs coordinate actin network formation within complex cellular environments.

## Methods

### Reagents

Unless otherwise specified, chemicals were obtained from Thermo Fisher Scientific (Waltham, MA). Cloning and SNAP-labeling reagents were obtained from New England Biolabs (Ipswich, MA).

### Plasmid construction

Mouse mDia1 (isoform 2; UniProtKB: O08808.1) constructs were either synthesized by GenScript (Piscataway, NJ) or subcloned from synthesized templates. For bacterial expression, mDia1 inserts were cloned into a modified pET23b vector containing an N-terminal Ulp1-cleavable 6×His-SUMO-tag, with AgeI and NotI restriction sites flanking mDia1 inserts. Constructs included: full-length (FL; residues 1-1255), constitutively active (CA; A256D, I259D), ΔDAD (1-1175), ΔARR (257-1255), NT (1-570), FH1-C (571-1255), and FH2-C (739-1255).

For mammalian cell expression, N-terminal SNAP-tagged constructs were cloned into pcDNA5/FRT. Inserts were transferred using AgeI/NotI restriction sites. FH1-FH2 (571-1175) was subcloned by GenScript and FH2 (739-1175) was generated by combining fragments from FH2-C and ΔDAD fragments.

Additional plasmids used in this study included GFP-β-actin (Addgene #31502; Watanabe and Mitchison, 2002), EMTB-2×mCherry (Addgene #26742; Miller and Bement, 2009), EB1-GFP (Addgene #17234; Piehl and Cassimeris, 2003) and GST-2×Myc-RhoA (Maiti et al., 2012; Nezami et al., 2006). Constructs not generated in-house were used as supplied.

### Protein expression and purification

6×His-SUMO-tagged mDia1 constructs were expressed in E. coli (Rosetta 2(DE3)-pRare2) and purified as described (Pimm et al., 2024) with the following modifications. Cells were lysed and bound to cobalt affinity resin (Cytiva, Marlborough, MA) in 1× PBS (140 mM NaCl, 2.5 mM KCl, 10 mM sodium phosphate dibasic, 1.8 mM sodium phosphate monobasic, pH 7.4) supplemented with 350 mM NaCl, 0.2 % Triton X-100, and 5 mM 2-mercaptoethanol and a protease inhibitor cocktail consisting of 0.5 µg/mL (each) antipain, leupeptin, pepstatin A, chymostatin, and aprotinin. ΔARR exhibited poor binding to cobalt resin and was instead purified using Ni-NTA (Qiagen, Germantown, MD) in poly prep columns (Bio-Rad, Hercules, CA) with modified buffer (1× PBS and 250 mM NaCl, without detergent or reducing agent) and eluted stepwise with 20-100% high-imidazole buffer (1× PBS, 250 mM NaCl, 400 mM imidazole, 0.05% NP-40, 5 mM 2-mercaptoethanol). After Ulp1-mediated cleavage of the 6×His-SUMO tag, all constructs were further purified by size-exclusion chromatography on a Superose6 Increase 10/300 column (Cytiva) equilibrated in HEKG10 buffer (50 mM HEPES (pH 7.4), 100 mM KCl, 10% glycerol, 0.02% NP-40, and 5 mM 2-mercaptoethanol). Peak fractions were pooled, concentrated using Amicon Ultra-4 50 kDa MWCO centrifugal filters (Millipore, Burlington, MA), flash frozen in liquid nitrogen, and stored at -80 ^o^C. At least three different batches of each mDia1 construct were purified and included in the in vitro analyses.

Human profilin-1 (PFN1) in a modified pET vector (pMW172) was expressed in E. coli (Rosetta 2(DE3)-pRare2) and purified by anion exchange chromatography on a HiTrapQ (Cytiva) equilibrated in 50 mM Tris-HCl (pH 8.0), followed by gel filtration on a Superdex75 increase 10/300 (Cytiva) in 50 mM Tris (pH 8.0), 50 mM KCl (Wolven et al., 2000; Pimm et al., 2022; Liu et al., 2022). GST-RhoA (2×Myc) in a modified pET vector was expressed in E. coli (Rosetta 2(DE3)-pRare2), purified on glutathione-Sepharose resin in a poly prep column (Bio-Rad), and eluted with a 0-500 nM concentration gradient of reduced glutathione, followed by gel filtration on a Superdex75 increase 10/300 (Cytiva) into 50 mM Tris pH 8.0, 100 mM NaCl, and 0.1 mM GTP, leaving the GST-tag intact (Li and Higgs, 2003; Maiti et al., 2012).

Rabbit skeletal muscle actin (RMA) was purified from acetone powder as described (Spudich and Watt, 1971; Pollard and Cooper, 1984). Briefly, powder stored at -80 ^o^C was rehydrated in G-buffer (3 mM Tris, pH 8.0, 0.5 mM DTT, 0.2 mM ATP, 0.1 mM CaCl_2_), clarified, polymerized overnight at 4 ^o^C, and pelleted. Pellets were dounce-homogenized, dialyzed against G-buffer for 48 h (two buffer exchanges), ultracentrifuged, and gel filtered (16/60 Superdex 200; Cytiva). RMA labeled on Cys374 with Alexa Fluor 488-labeled and pyrene (maleimide) or on lysine residues with Alexa Fluor 488 (NHS ester) were generated as described (Cooper et al., 1983; Kuhn and Pollard, 2005) or lysine residues (NHS ester) (Hertzog and Carlier, 2005; Pimm et al., 2022, 2024). Briefly, for Cys374 labels, actin was dialyzed into DTT-free G-buffer, polymerized (1 mg/mL), and labeled overnight with 7-10-fold molar excess Alexa Fluor 488- or pyrene-maleimide in 25 mM imidazole (pH 7.5), 100 mM KCl, 0.15 mM ATP, and 2 mM MgCl_2_. Filaments were pelleted, dialyzed, and gel filtered as above. For lysine labeling, actin was dialyzed into HEPES-based G-buffer (3 mM HEPES, pH 8.2, 0.5 mM DTT, 0.3 mM ATP, 0.1 mM CaCl_2_), polymerized, and labeled on filaments in 3 mM HEPES (pH 8.2), 0.5 mM DTT, 0.3 mM ATP, 1 mM MgCl_2_, and 50 mM KCl prior to pelleting and gel filtration.

Protein concentrations and purity were determined by Coomassie-stained SDS-PAGE densitometry relative to a BSA standard curve. Concentrations of labeled actin were calculated spectrophotometrically using ε290 = 25,974 M^-1^ cm^-1^ for actin and ε495 = 71,000 M^-1^ cm^-1^ for Alexa Fluor 488, applying a correction factor of 0.11. Labeling efficiency was determined from corrected absorbance ratios.

### Pyrene actin polymerization assays

Bulk actin assembly was monitored using pyrene fluorescence. Reactions contained 2 µM Mg-ATP actin (5% pyrene-labeled) supplemented with 1 nM mDia1 constructs in the presence or absence of 5 µM profilin-1. Actin was exchanged with 1 mM MgCl_2_ and 10 mM EGTA for 2 min before polymerization was initiated by the addition of 20× initiation mix (final reaction concentration of 2 mM MgCl_2_, 0.5 mM ATP, and 50 mM KCl). For each experiment, reactions were assembled in parallel and initiated by rapid addition of actin using a multichannel pipette to ensure synchronized start times. Fluorescence (excitation 365 nm, emission 407 nm) was recorded continuously in a plate reader. Three technical replicates were averaged per condition. Assays shown in Figures 1 and 2 were performed on the same instrument during the same experimental session but are presented separately for clarity.

### In vitro actin assembly assays via TIRF microscopy

Flow chambers were constructed using biotin-PEG-coated #1.5 coverslips (24 × 60 mm) adhered to µ-slide VI chambers (IBIDI, Fitchburg, WI) using 0.12 mm SA-S-Secure Seal double-sided tape (Grace Bio Labs, Bend, OR) and 5-min epoxy (Loctite/Henkel Corp., Rocky Hill, CT), as described (Henty-Ridilla, 2022; Haarer et al., 2023). Chambers were sequentially incubated with 1% BSA, 0.005 mg/mL streptavidin, 1% BSA, and 1× TIRF buffer (20 mM imidazole [pH 7.4] 50 mM KCl, 1 mM MgCl_2_, 1 mM EGTA, 0.2 mM ATP, 10 mM DTT, 40 mM glucose, 0.25% methylcellulose [4000 cP], 10 mg/mL glucose oxidase, and 2 mg/mL catalase) prior to introduction of actin polymerization reactions. Unless otherwise noted, reactions contained 1 µM total rabbit skeletal muscle actin (10% Alexa Fluor 488-labeled; 6 nM biotin-actin), with or without 1 nM mDia1 and/or 5 µM profilin-1. For activation experiments with full-length mDia1 or ΔARR, 0.5 µM GST-RhoA was included.

For figure panels using lysine-labeled actin (Figures 1D, 2C, and S1D-E), imaging was performed on a Leica DMi8 microscope equipped with a 100× Plan Apo 1.47 NA oil-immersion objective, 488 nm solid-state laser (120 mW laser set to 5% power, 100 ms exposure), and an iXon Life 897 EMCCD camera (Andor, Belfast, UK), yielding a (81.9 µm)^2^ field of view (FOV). Time-lapse images were acquired every 5 s for 16-20 min using LAS X software (Leica Microsystems, Wetzlar, Germany).

For experiments and figure panels using Cys374-labeled actin (Figures 1E-G, 2D-F, and S1F-G), imaging was performed on a Nikon iLAS TIRF microscope equipped with a 60× 1.49 NA oil-immersion objective, 488 nm excitation (90 mW laser set to 10% power; 100 ms exposure) (Nikon, Inc, Melville, NY), and a Prime BSI express sCMOS camera (Teledyne Photometrics, Tucson, AZ), yielding a (111.41 µm)^2^ field of view. Time-lapse images were captured every 5 s for 15 min. Raw images were minimally processed in FIJI for background subtraction. These adjustments were applied uniformly across conditions within each experiment.

Images were exported as TIFF files and analyzed in FIJI (Schindelin et al., 2012). Filament nucleation was quantified as the number of filaments per field of view (FOV) at 100 s after reaction initiation. Elongation rates were determined by measuring filament length at 5-7 time points per filament, and rates were calculated from the slope of these measurements. Slopes were converted to subunits per micron per s using a conversion factor of 370 subunits per micron of filament (Pollard et al., 2000). For profilin-mediated fast formin elongation, rates were measured only from dim formin-associated filaments. The percentage of dim filaments in Cys374-labeled actin reactions (Supplementary Figure 1F-G) was calculated as the fraction of dim filaments relative to total filaments at 200 s.

### Mammalian cell culture and transfection

Mouse Neuroblastoma-2A (N2A) cells (ATCC, Manassas, VA) were maintained in DMEM supplemented with 10% FBS (GenClone; Genesee Scientific, El Cajon, CA), 2 mM L-glutamine, and penicillin (50 U/mL)-streptomycin (50 µg/mL) at 37 ^o^C in 5% CO_2_. Clonal mDia1 knockout (mDia1-KO) lines were derived from pooled CRISPR-edited N2A cells (Synthego, Menlo Park, CA) and validated by immunoblotting (Supplementary Figure 2). For live-cell imaging experiments, cells were seeded onto 9.6 cm^2^ glass-bottom dishes (MatTek Corp., Ashland, MA) at 15,000-25,000 cells per dish and transfected the following day using Lipofectamine 3000. SNAP-mDia1 constructs were used at 250 ng per dish. Where indicated, GFP-actin, EMTB-2×mCherry, or EB1-GFP were transfected at 100 ng each. Cells were imaged 24-48 h post-transfection.

### SNAP labeling and live cell imaging

For live-cell imaging, growth medium was replaced with FluoroBrite DMEM supplemented with 20 mM HEPES (pH 7.4), 10% FBS, and 2 mM L-glutamine. SNAP-tagged proteins were labeled with 75 nM SNAP-Cell 647-SiR ligand for 20 min prior to imaging.

Images were acquired on a Nikon Ti2-E spinning disk confocal SoRa microscope (Nikon Instruments) equipped with 405-, 488-, 561-, and 640-nm lasers, a CSU-W1 scan head (Yokogawa, Ishikawa, Japan), a Plan Apo 60× 1.4 NA oil-immersion objective, and either a Prime BSI sCMOS (Teledyne Photometrics, Tucson, AZ) or ORCA-Quest 2 qCMOS camera (Hamamatsu Photonics, Hamamatsu City, Japan). Cells were maintained at 37 ^o^C in a stage-top environmental chamber (OKOlab, Sewickley, NJ).

Time-lapse TIRF imaging was performed on the Nikon iLAS system (described above) with cells maintained in a stagetop incubator (OKOlab). Images were acquired every 1 s with 100 ms exposure for 2 min using 488 nm excitation. SNAP-mDia1-expressing cells were verified for expression; however, only the EB1-GFP channel was analyzed. Mock-transfected cells lacking SNAP signal were included as controls. Microtubule dynamics were quantified by tracking EB1-GFP comets using plusTipTracker software (version 2.2.0; Applegate et al., 2011) in MATLAB (version R2024b; The MathWorks, Natick, MA). Comet detection and velocity measurements were performed on the original time-lapse image sequences. For visualization only, EB1-GFP dynamics were also displayed as maximum-intensity projections over short temporal windows (10 s), which illustrate comet trajectories. Each data point represents either the number of comets per cell or mean comet velocity, calculated over the 2 min acquisition period. For experiments comparing puncta intensity (e.g., bright versus dim), laser power, exposure time, camera gain, and binning were held constant within each experiment. Unless otherwise indicated, time-lapse imaging was performed at 1-5 s intervals.

### Filopodia quantification and analysis

Filopodia number and length were quantified from maximum-intensity projections of SoRa Z-stacks using FIJI. Lengths were measured manually along the filopodial axis from base to tip from Z-stacks (0.3 µm intervals) encompassing the entire cell. Each point represents an individual filopodium (n = 22-40 per condition for WT and n = 17-28 per condition for knockout cells) from multiple cells across N = 2-3 independent transfections (indicated by shading). Filopodia elongation rates were determined from time-lapse movies by measuring changes in filopodial length over time from maximum-intensity projections generated from three Z-sections at 0.3 µm intervals. Each data point represents the mean elongation rate per cell, averaged across all filopodia analyzed (n = 12-15 filopodia per condition across multiple cells and transfections, indicated by shading).

For SNAP-mDia1 puncta analysis, temporal dynamics were visualized by projecting the first ten frames of the SNAP-mDia1(CA) channel as a time-encoded image, where frame progression is represented by color intensity; more color variation indicates greater dynamics, whereas white indicates static signal over the ten frame window. Puncta movement was further analyzed using kymographs generated along filopodial axes. Line scans were drawn from the filopodial tip toward the base, with the tip defined as the distal 1.5 µm and the shaft defined as 1.5-3 µm from the tip (as indicated by the red dotted line in Figure 4H). SNAP-mDia1(CA) maximum pixel intensity was quantified from these scans, and puncta intensity in the tip and shaft regions was compared. Each data point represents puncta from individual filopodia across multiple cells.

### Immunoblotting

Expression of SNAP-tagged mDia1 proteins in N2A cells was assessed 24-48 h post-transfection. Cells were washed with PBS and lysed directly in SDS sample buffer. Lysates were heated to 95 ^o^C for 5 min, sonicated briefly, clarified by centrifugation at 21,000 × *g* for 5 min, and 10-15 µL was loaded per lane on polyacrylamide gels. When indicated, known amounts of purified mDia1 were included as standards. Proteins were separated by SDS-PAGE and transferred to nitrocellulose membranes.

Membranes were blocked in PBS containing 5% BSA and washed in PBST (1×PBS and 0.1% Tween-20). Primary antibodies were applied in PBST + 5% BSA at room temperature for 2 h or at 4 ^o^C overnight, including: rabbit anti-Diaph1 (Abcam, #ab129167; 1:1000), mouse anti-Dia1 (Santa Cruz, #sc-373807; 1:500), mouse anti-β-tubulin (DSHB, #E7; 1:2000), mouse anti-GAPDH (Proteintech, #60004-1-lg; 1:50,000), and rabbit anti-α-tubulin (Abcam, #ab18251; 1:3000). Blots were washed and probed with fluorescent secondary antibodies (LI-COR IR680 or IR800, 1:5000-1:10,000) in PBST and 5% BSA, then imaged using a LI-COR Odyssey system.

### Statistical analysis, software, and data availability

Data are presented as mean ± SEM unless otherwise indicated with details of sample size and tests provided in figure legends. Statistical analyses were performed using GraphPad Prism 10 (version 10.4.2). Tests and replicate numbers are provided in figure legends. One-way ANOVA with Tukey’s post hoc or student’s t-tests were used for all comparisons. Statistical significance was defined as α = 0.05. Molecular models and visualizations were prepared using PyMOL (version 3.1.8). Figures were prepared using Adobe Illustrator 2026 (version 30.3). Source datasets are available via Zenodo: 10.5281/zenodo.20404275.

## Supplemental information

Supplemental information includes 4 figures and 5 movies.

## Acknowledgements

We are grateful to Marc Ridilla and David Amberg for helpful discussions and comments on this manuscript. We also thank M. Groening for readily inspiring exploration of FH2-shaped structures. This work was supported by National Institutes of Health grant R35GM133485 to JLH-R.

## Author contributions

**Brian K. Haarer**: Data curation, formal analysis, investigation, methodology, validation, writing (original draft, reviewing, and editing). **Amanda Davenport**: Formal analysis, investigation, writing (reviewing and editing). **Laura M. Haney**: Formal analysis, investigation, writing (reviewing and editing). **Morgan L. Pimm**: investigation, validation, writing (reviewing and editing). **Alexander D. Nobles**: investigation, writing (reviewing and editing). **Krishna Patel**: investigation, writing (reviewing and editing). **Jessica L. Henty-Ridilla**: Conceptualization, data curation, formal analysis, investigation, project administration, supervision, resources, funding acquisition, validation, visualization, writing (original draft, reviewing, and editing).

## Supplemental Information

**Figure S1.**
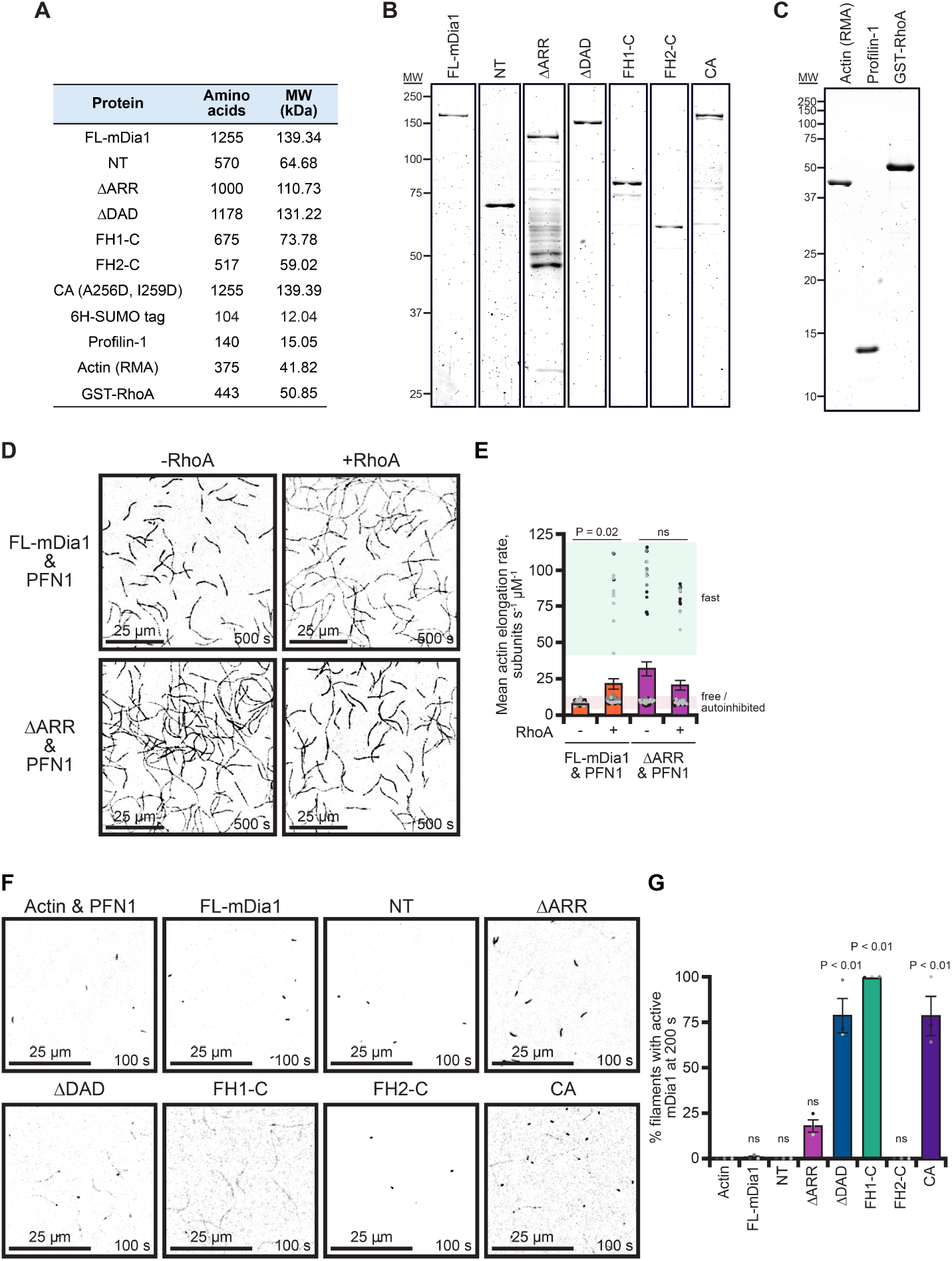
Protein purity, RhoA-dependent activation, and processivity of mDia1 constructs. **(A)** Table summarizing the predicted molecular weights of mDia1 constructs used in this study. **(B)** Coomassie-stained SDS-PAGE gel of purified mDia1 proteins. Approximately 100 ng of each protein was loaded to assess purity. **(C)** Coomassie-stained SDS-PAGE gel of additional purified proteins used in this study: rabbit skeletal muscle actin, profilin-1 (PFN1), and GST-RhoA. Approximately 300 ng of each protein was loaded. **(D)** Representative TIRF images of actin assembly reactions containing 1 µM actin (10% Alexa Fluor 488-labeled on lysine), 5 µM PFN1, and 1 nM of the indicated mDia1 construct, in the absence or presence of 0.5 µM GST-RhoA. Images were acquired 500 s after initiation of polymerization. Scale bars, 25 µm. **(E)** Actin filament elongation rates from reactions as in (D). Each dot represents an individual filament (n = 68 filaments per condition from N = 3 independent reactions; shading indicates independent experiments). Statistical analysis was performed by one-way t-tests, comparing to RhoA-negative control; ns, not significant (P > 0.05). **(F)** Representative TIRF images of reactions containing 1 µM actin (10% Alexa Fluor 488-labeled on Cys374), 5 µM PFN1, and 1 nM of the indicated mDia1 construct. Under these conditions, processive elongating formin-associated filaments appear dimmer than non-formin-associated filaments due to profilin-mediated suppression of filament brightness (Kovar et al., 2006). Images were acquired 100 s after polymerization initiation. Scale bars, 25 µm. **(G)** Quantification of the fraction of dim filaments per field of view. The percentage of dim filaments serves as an operational readout of processive formin engagement under these assay conditions. Each dot represents the mean of 3 fields of view per independent reaction (N = 3; shading indicates independent measurements). Statistical analysis was performed by one-way ANOVA, compared to actin and PFN1 control; ns, not significant (P > 0.05).

**Figure S2.**
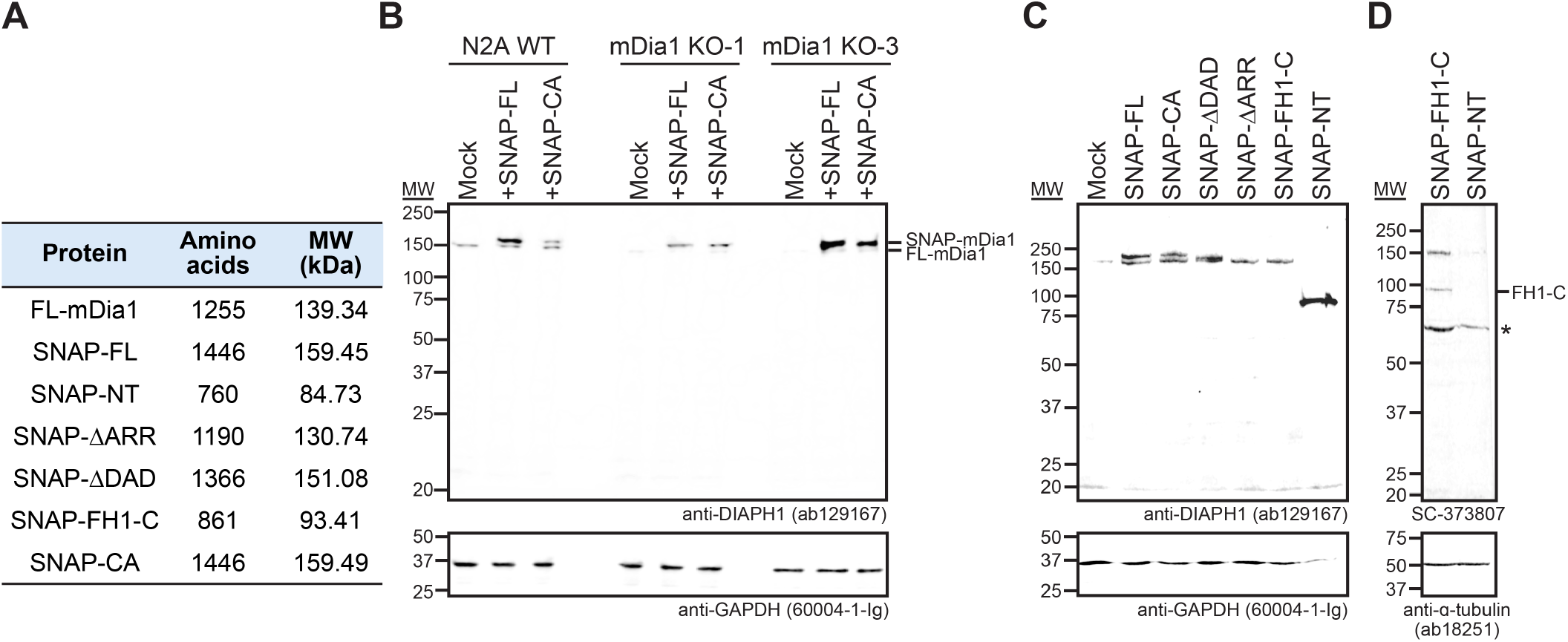
Validation of mDia1 knockout lines and SNAP-mDia1 plasmid expression via Western blots. **(A)** Table summarizing the predicted molecular weights of untagged full-length (FL-)mDia1 and SNAP-tagged mDia1 constructs used in cells. **(B)** Western blot analysis of mDia1 protein levels in wild-type N2A cells and two independent mDia1 knockout (KO) lines transfected with mock, SNAP-FL-mDia1, or SNAP-mDia1(CA). Blots were probed with rabbit anti-DIAPH1 (Abcam ab129167; 1:1000; N-terminal epitope) and mouse anti-GAPDH (ProteinTech 60004-1-lg; 1:50,000) as a loading control. **(C)** Western blot of wild-type N2A cells expressing the indicated SNAP-mDia1 constructs. Blots were probed with rabbit anti-DIAPH1 (Abcam ab129167; 1:1000; N-terminal epitope) and mouse anti-GAPDH (ProteinTech 60004-1-lg; 1:50,000) as a loading control. **(D)** Western blot of cells expressing the indicated constructs probed with mouse anti-DIAPH1 (Santa Cruz sc-373807; 1:500; C-terminal epitope) and rabbit anti-α-tubulin (Abcam ab18251; 1:3000) as a loading control. Asterisk denotes a non-specific band. N- and C-terminal antibodies were used because epitope accessibility differs across constructs, limiting detection of certain truncation mutants with a single antibody.

**Figure S3.**
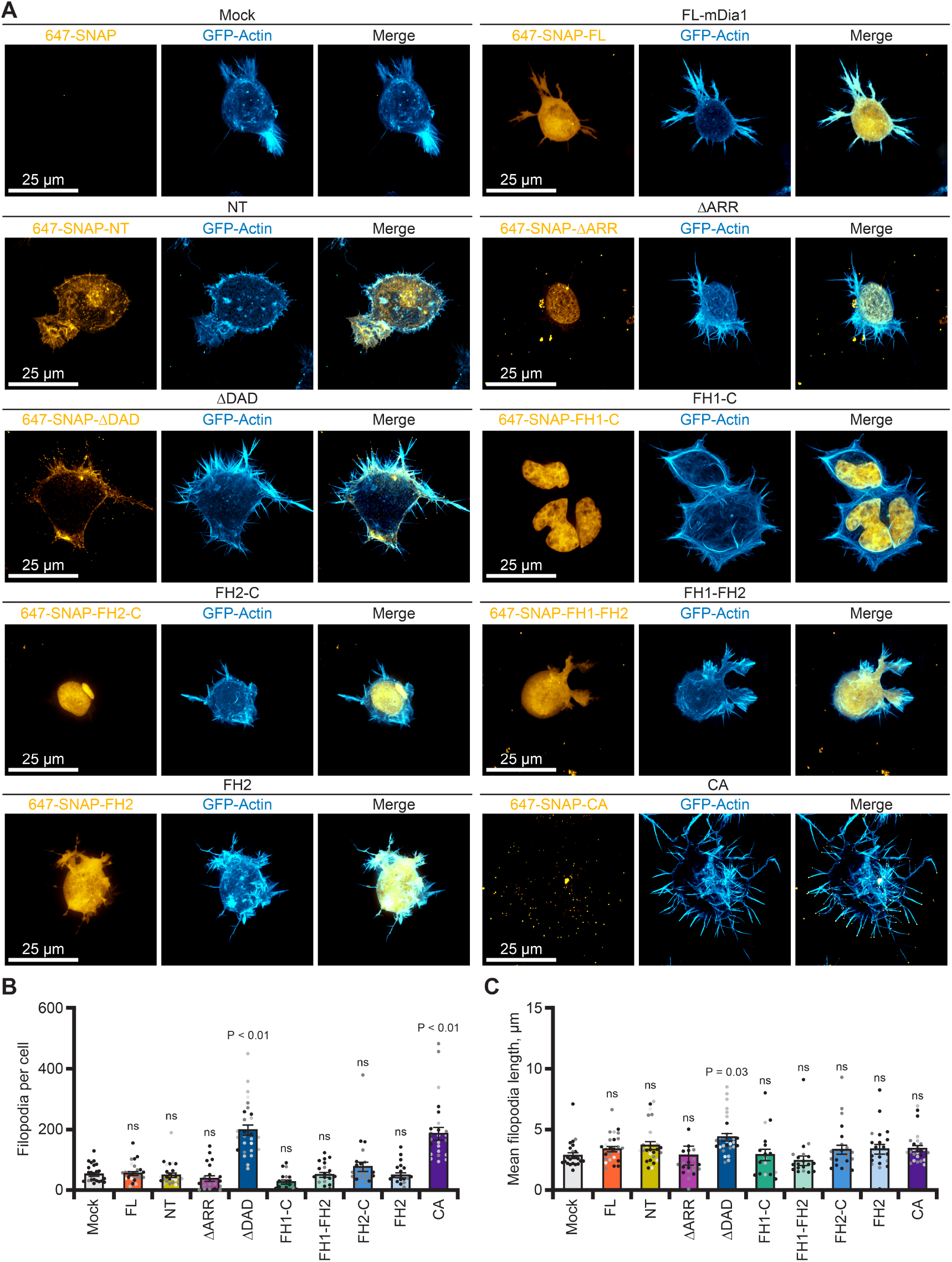
SNAP-mDia1 constructs in mDia1 knockout N2A cells. **(A)** Live-cell confocal images of mDia1 knockout (KO) N2A cells expressing the indicated N-terminally SNAP-tagged mDia1 constructs. SNAP signal (gold), GFP-actin (blue), and merged images are shown as maximum intensity projections. Scale bars, 25 µm. **(B)** Quantification of filopodia number per cell from 3D image stacks as in (A). Each dot represents a single cell (n = 17-28 cells per condition); shading denotes N = 2-3 independent experiments (separate transfections performed on different days). Statistical analysis was performed by one-way ANOVA, compared to mock; ns, not significant (P > 0.05). **(C)** Mean filopodial length per cell from datasets as in (A). Each dot represents the average filopodial length for a single cell (n = 17-28 cells per condition). Shading denotes N = 2-3 independent experiments. Statistical analysis was performed by one-way ANOVA, compared to mock; ns, not significant (P > 0.05). Expression of activated SNAP-mDia1 constructs in mDia1 KO cells recapitulated the puncta formation and filopodia phenotypes observed in wild-type cells, indicating that these effects are intrinsic to the expressed constructs and are not altered by loss of endogenous mDia1.

**Figure S4.**
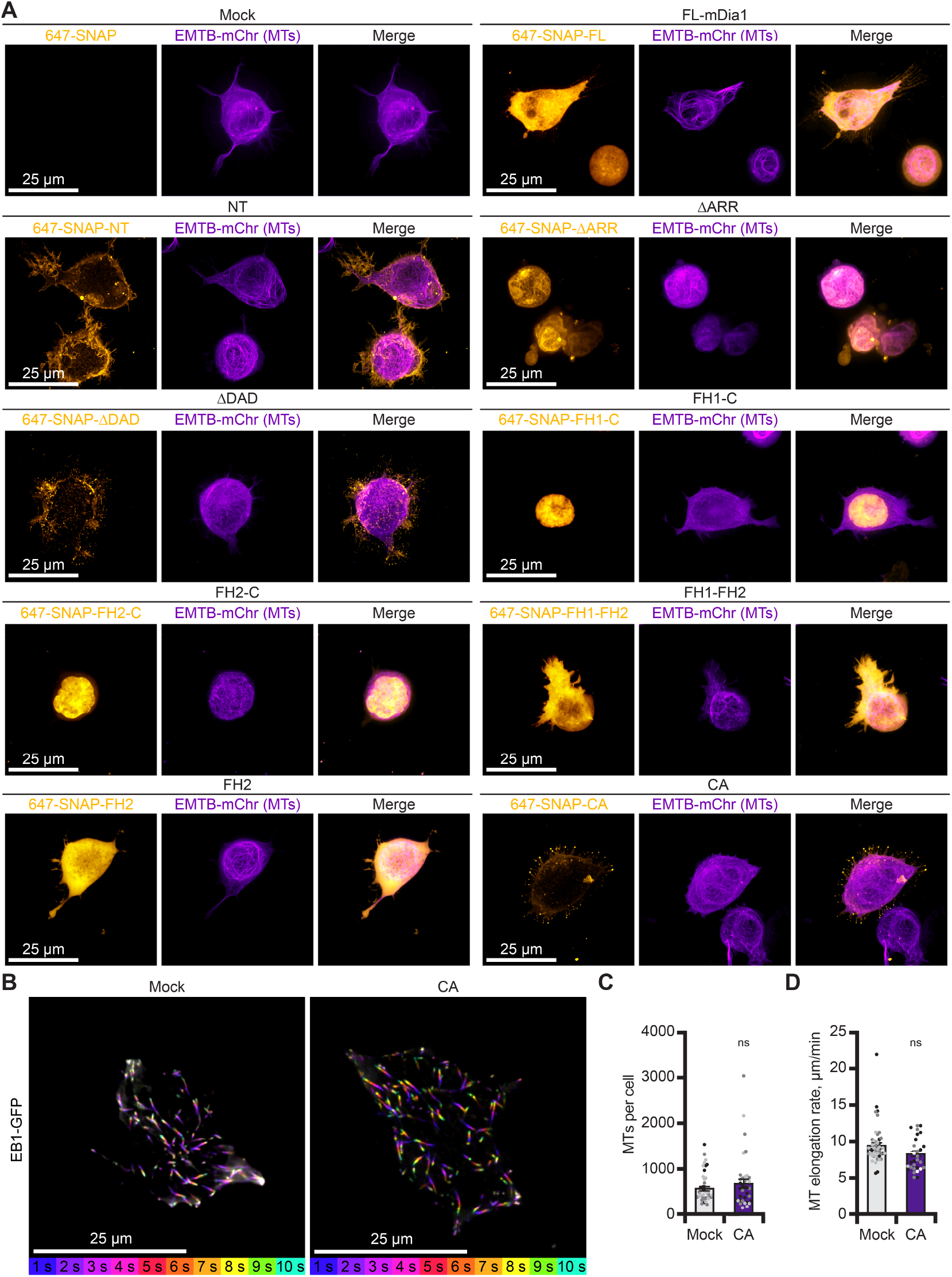
Microtubule dynamics are not altered by SNAP-mDia1(CA) expression in N2A cells. **(A)** Live-cell confocal images of wild-type N2A cells expressing the indicated N-terminally SNAP-tagged mDia1 constructs. SNAP signal (gold), EMTB-2×mCherry (purple), and merged images are shown as maximum intensity projections. Scale bars, 25 µm. These images correspond to the cells shown in Figure 3A and include the microtubule mark to assess potential spatial association. Under these conditions, SNAP-mDia1 constructs do not show detectable enrichment along microtubules. **(B)** Representative time-encoded projection of time-lapse TIRF imaging of EB1-GFP comets over 10 s. Frames are color-coded by time; greater color variation indicates dynamic signal, whereas white indicates minimal or no movement over the acquisition period. Scale bars, 25 µm. **(C)** Microtubule count per cell quantified from EB1-GFP comets using plusTipTracker. Each dot represents the total number of comets per cell. Statistical analysis was performed using a one-way t-test, compared to mock; ns, not significant (P > 0.05). **(D)** Microtubule growth rates measured from EB1-GFP comets using plusTipTracker. Each dot represents the mean microtubule growth rate per cell over 2 min acquisition period (images acquired every 1 s). Shading indicates N = 3 independent experiments. Statistical analysis was performed as in (C).

**Movie 1. Autoinhibition restrains mDia1-driven actin assembly**. Time-lapse TIRF movie of 1 µM actin (10% Alexa 488-Lys-label) polymerizing with 1 nM indicated mDia1 constructs. Images acquired every 5 s. Playback, 10 frames per s (FPS). Scale bar, 25 µm.

**Movie 2. Profilin accelerates elongation by activated mDia1**. Time-lapse TIRF movie of 1 µM actin (10% Alexa 488-Lys-label) polymerizing with 1 nM indicated mDia1 constructs in the presence of 5 µM profilin-1. Images acquired every 5 s. Playback 10 FPS. Scale bar, 25 µm.

**Movie 3. Bright and dim filament populations during formin-mediated growth**. Time-lapse TIRF movie of 1 µM actin (10% Alexa 488 Cys374-label) polymerizing with 1 nM indicated mDia1 constructs and 5 µM profilin-1. Bright and dim filaments are visible. Images acquired every 5 s. Playback 10 FPS. Scale bar, 25 µm.

**Movie 4. SNAP-mDia1(CA) puncta in filopodia**. Live-cell SoRa confocal imaging of SNAP-mDia1(CA) (gold) and GFP-actin (blue) in an N2A cell corresponding with Figure 4A-B. Bright tip-localized puncta and dim shaft puncta are visible within filopodia. Images acquired every 4 s. Playback 10 FPS. Scale bars, 10 µm and 2 µm (inset).

**Movie 5. SNAP-mDia1(CA) puncta in filopodia imaged by TIRF microscopy**. Live-cell TIRF imaging of SNAP-mDia1(CA) (gold) and GFP-actin (blue) in an N2A cell corresponding with Figure 4D-E. An actively growing filopodium has an increasing SNAP-mDia1(CA) signal (arrow). Images acquired every 1 s. Playback 10 FPS. Scale bars, 10 µm and 2 µm (inset).

